# Grazing of the free-living filamentous brown seaweed, *Pilayella littoralis*, by the amphipod *Gammarus tigrinis*

**DOI:** 10.1101/2023.05.23.541919

**Authors:** Steven L. Miller, Robert T. Wilce

## Abstract

*Pilayella littoralis* is a common brown alga with a cosmopolitan distribution. A distinct form of the alga is found free-living in Nahant Bay, Massachusetts. Characteristics that distinguish the free-living form include radial branching that results in a ball-like morphology, reproduction by fragmentation of filaments rather than by spores, and year-round persistence of the population. The amphipod *Gammarus tigrinis* is frequently found within the floating drifts of *P. littoralis*. We conducted gut content analyses to document that *G. tigrinis* consumes *P. littoralis*. Culture studies revealed that ingested *P. littoralis* could survive and grow from fecal pellets produced by *G. tigrinis*. In addition, inefficient grazing produces vegetative fragments (propagules) that can increase numbers within the free-living population. δ^13^C values for the amphipods and *P. littoralis* averaged -17.4 o/oo and -17.7 o/oo, respectively, suggesting that *G. tigrinis* acquires most of its carbon from *P. littoralis*.

## Introduction

The effects of grazers on the abundance, distribution, and morphology of seaweeds are well documented (Underwood 1979, Slocum 1980, Lubchenco and Gaines 1981, Gaines and Lubchenco 1982, Hawkins and Hartnoll 1983). Herbivores may also facilitate the reproduction and dispersal of seaweeds when grazed filaments are released as drift or when seaweeds survive ingestion and germinate from fecal pellets (Santelices et al. 1983, Santelices and Correa 1985, Breeman and Hoeksema 1987, Santelices and Ugarte 1987). In freshwater, changes in community structure can result from the differential ability of freshwater algae to survive ingestion (Gibor 1956, Porter 1973, 1976, Nicotri 1977). A wide variety of seaweeds also survive ingestion by herbivores, including sea urchins, many gastropods, and a chiton (Santelices et al. 1983, Santelices and Correa 1985, Breeman and Hoeksema 1987, Santelices and Ugarte 1987).

We investigated the relationship between the filamentous brown seaweed *Pilayella littoralis* (L.) Kjellman and the amphipod, *Gammarus tigrinis* Sexton, in Nahant Bay, Massachusetts, where *P. littoralis* occurs as a free-living perennial population (Wilce et al. 1982, Miller 1988, Pregnall and Miller, 1988). The alga reproduces solely by vegetative fragmentation. Reproductive unilocular and plurilocular organs are rarely present and do not contribute significantly to population growth (Wilce et al. 1982). Nuisance accumulations of *P. littoralis* occur in the surf zone of beaches fringing Nahant Bay and often reach densities of 0.85 g dry weight/liter. *G. tigrinis* is present year-round and accounts for at least 15 percent of the dry weight of the free-living *P. littoralis* community during summer months (pers. comm. T. Briggs).

To evaluate the effects of *Gammarus tigrinis* grazing on the free-living *Pilayella littoralis* population in Nahant Bay, Massachusetts, we conducted gut analyses to establish if *G. tigrinis* consumes *P. littoralis in situ*. After we observed algal filaments growing out of amphipod fecal pellets in culture, we conducted laboratory tests to describe and quantify the phenomenon. After confirming that gut contents of *G. tigrinis* collected *in situ* mainly contained *P. littoralis*, we performed stable carbon isotope analyses to evaluate if *P. littoralis* was a primary dietary carbon source for *G. tigrinis*.

## Methods

We analyzed gut contents from samples of the amphipod *Gammarus tigrinis*, collected periodically with free-living *Pilayella littoralis* as part of a phenological study of the alga in Nahant Bay, Massachusetts (Figure 1). Ingested materials were identified using a compound microscope.

**Figure 1.**
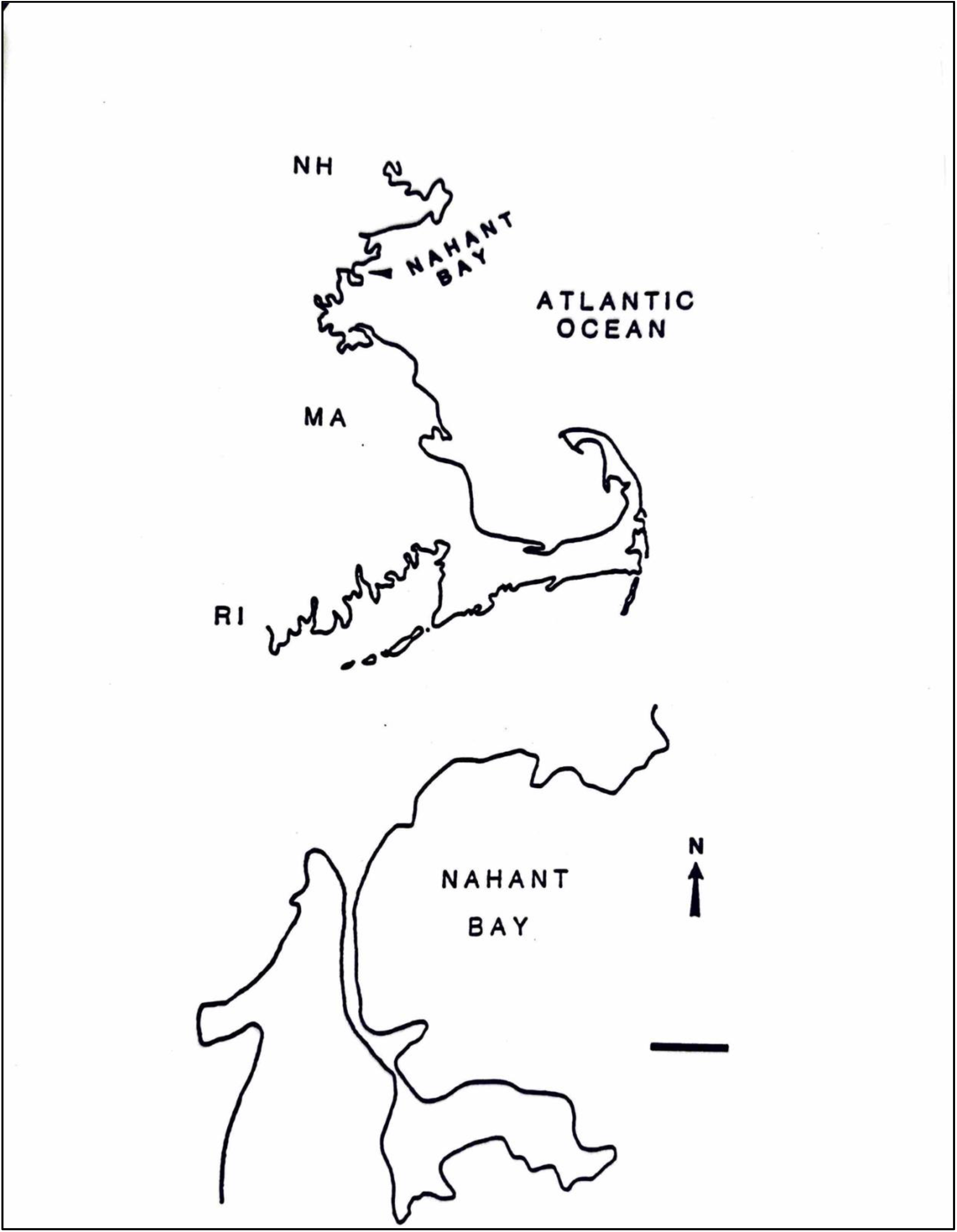
Map of Massachusetts coastline and Nahant Bay. Bar = 1 Km.

We collected *Gammarus tigrinis* from *in situ* populations of free-living *Pilayella littoralis* and collected fecal pellets produced after one and twenty-four hours. The amphipods were first rinsed three times in sterile seawater. An inverted microscope was used to observe the fecal pellets to ensure no contaminants were present. Fecal pellets were rinsed once in sterile seawater and then set aside for incubation at 10° C and 30 uE/m^2^/sec under long days (16:8 light:dark cycle). As a control, seawater used to rinse the pellets was checked for subsequent algal development. After one week, pellets were transferred to either 10 ml of sterile F/4 culture medium (modified after Guillard and Ryther 1962) in a plastic petri dish or 200 ml of F/4 media in a glass storage dish. Cultures were transferred to new media weekly and were monitored for 30 days to check for the development of algal filaments from fecal pellets. Fecal pellets were also collected from amphipods placed in unialgal cultures of free-living *P. littoralis*.

Stable carbon isotope analyses (δ^13^C) were performed on amphipods collected *in situ* and starved for one week to evacuate their digestive tract. In addition, the δ^13^C value of free-living *Pilayella littoralis* was determined from material collected with the amphipods and meticulously picked clean of other algae and invertebrates (mostly amphipods and copepods). In addition, we obtained stable carbon isotope values for the free-living *P. littoralis* community, including unsorted algae, animals, and debris. We tested the null hypothesis that no difference in carbon isotope ratios exists between free-living *P. littoralis* and *Gammarus tigrinis*. Rejection of the null hypothesis (t-test p ≤ 0.05) implies that the amphipods assimilate considerable carbon from a source other than *P. littoralis*.

Samples were prepared for stable carbon isotope analysis following Dunton and Schell (1987). Isotope measurements were performed on a Micromass 602E dual inlet, double collector isotope-ration mass spectrometer. Results were expressed as δ^13^C values relative to the limestone standard PDB (Craig 1987), where:

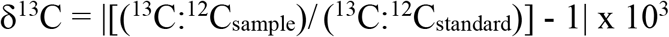

Secondary standards were run daily as cross-calibration checks of the AER oil working standard, combusted daily. NBS-22 (δ^13^C = -29.8 o/oo) was used as a common secondary standard.

## Results

Analyses of *Gammarus tigrinis* gut contents revealed *Pilayella littoralis* comprised much of the material ingested by these animals, though unidentified digested material that resembled *P. littoralis* was common along with other filamentous algae, diatoms, and debris (Figures 2, 3). In addition, cell wall ghosts with cellular dimensions similar to *P. littoralis* could frequently be discerned among the debris. Two other macroalgae infrequently grew from fecal pellets produced by amphipods collected in the field, including *Enteromorpha* sp. and an identified brown alga. The survivorship of ingested filaments was low, with three pellets out of 80 producing live algal filaments. *P. littoralis* was the only alga that grew vigorously from the fecal pellets (Figure 4). No macroalgae grew in the sterile seawater rinses of amphipods or fecal pellets, although numerous diatoms and ciliates eventually dominated these cultures. Filaments of *P. littoralis* also grew from fecal pellets produced by amphipods fed unialgal *P. littoralis* (Figure 5).

**Figure 2.**
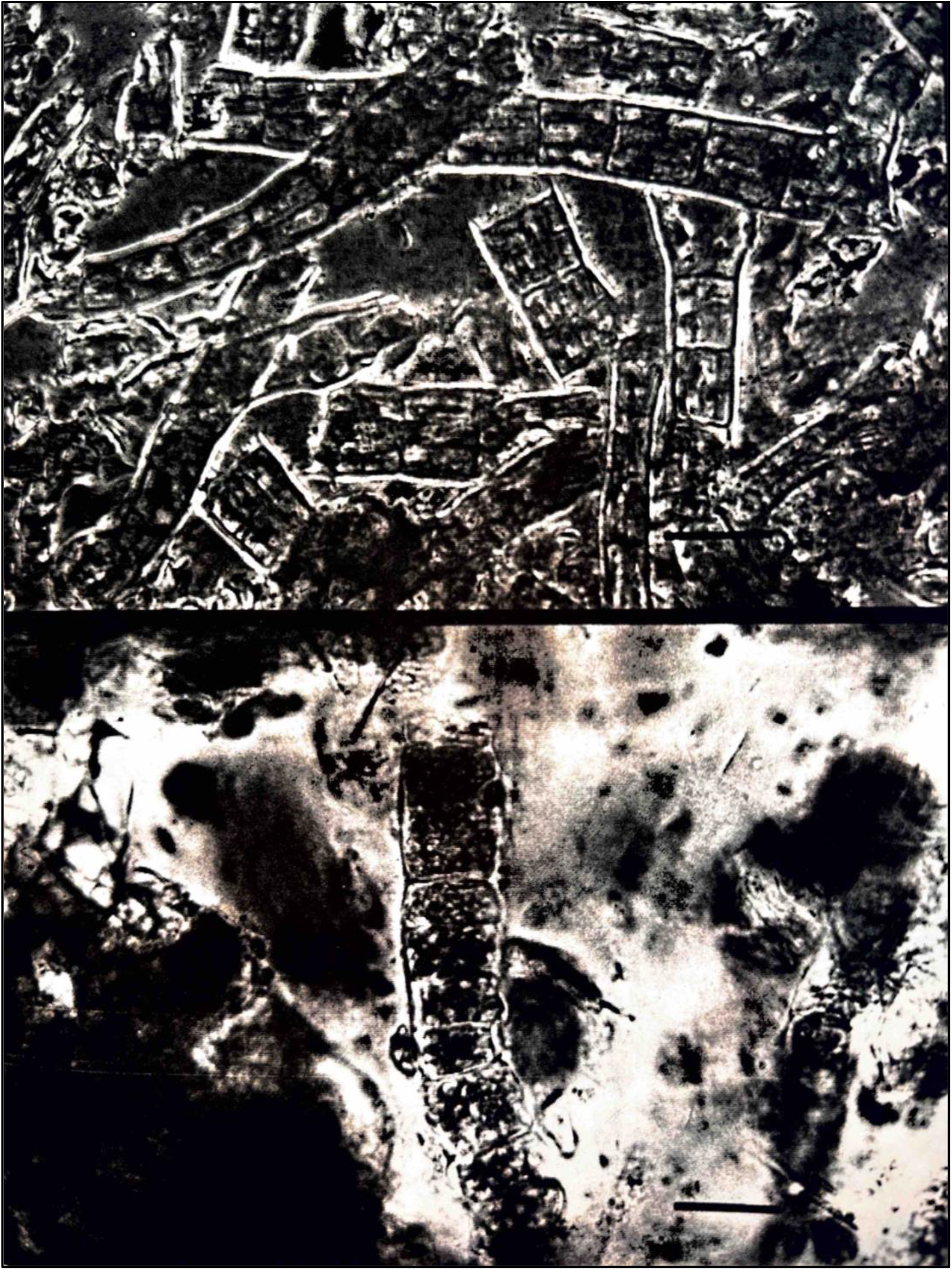
Gut content of *Gammarus tigrinis* collected among free-living *Pilayella littoralis*: brown algal filaments. Top; gut with algal filaments that resemble *Sphacelaria* sp., note pluriseriate condition. Bottom; intact filament resembling *P. littoralis*. Scale bars = 25 um.

**Figure 3.**
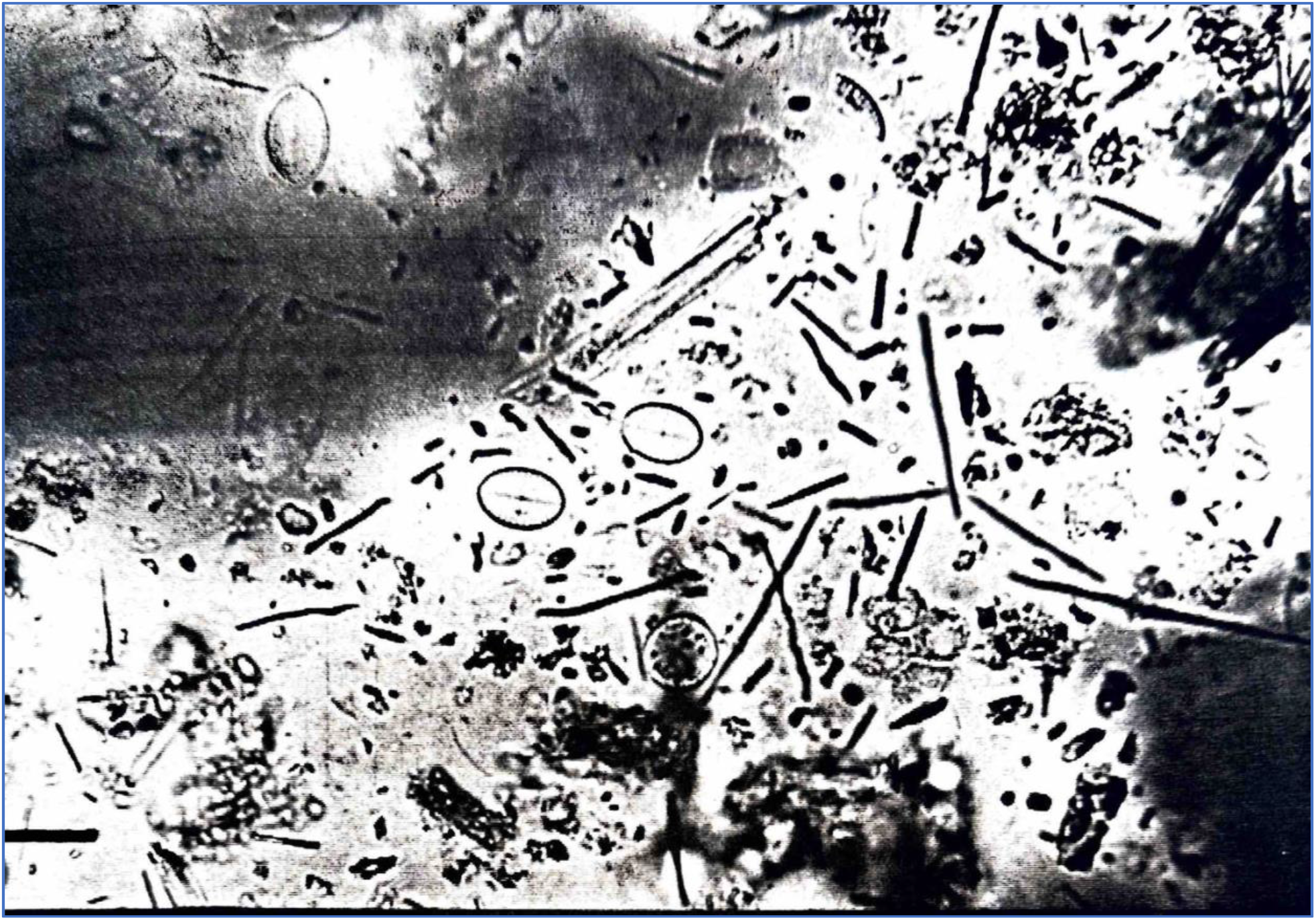
Gut content of *Gammarus tigrinis* collected among free-living *Pilayella littoralis*: diatoms, debris, and spicule-like material. Scale bar = 10 um.

**Figure 4.**
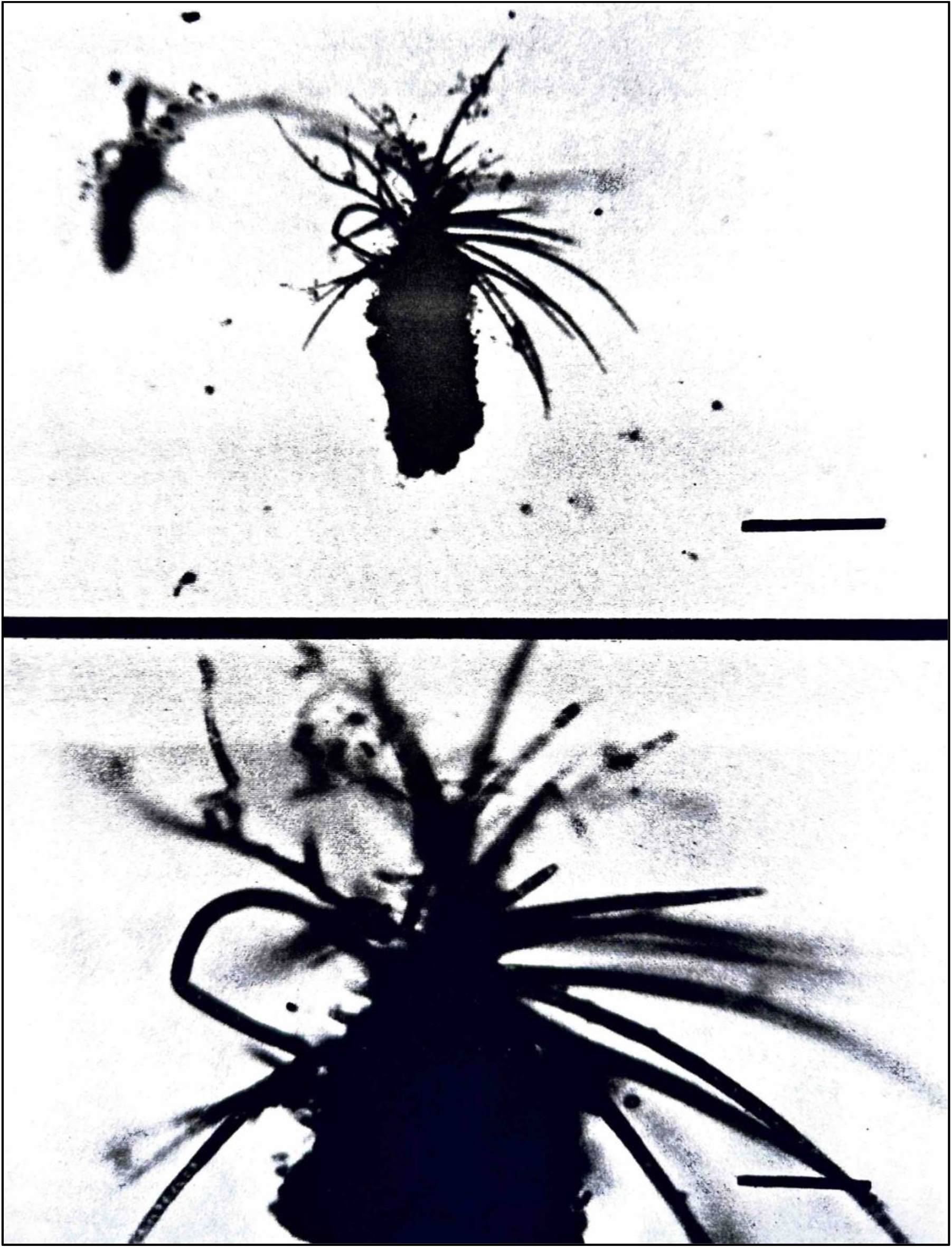
Fecal pellet produced by *Gammarus tigrinis* collected from within a cloud of free-living *Pilayella littoralis*. See text for germination conditions and sampling procedure. Scale bars; top = 400 um, bottom = 125 um.

**Figure 5.**
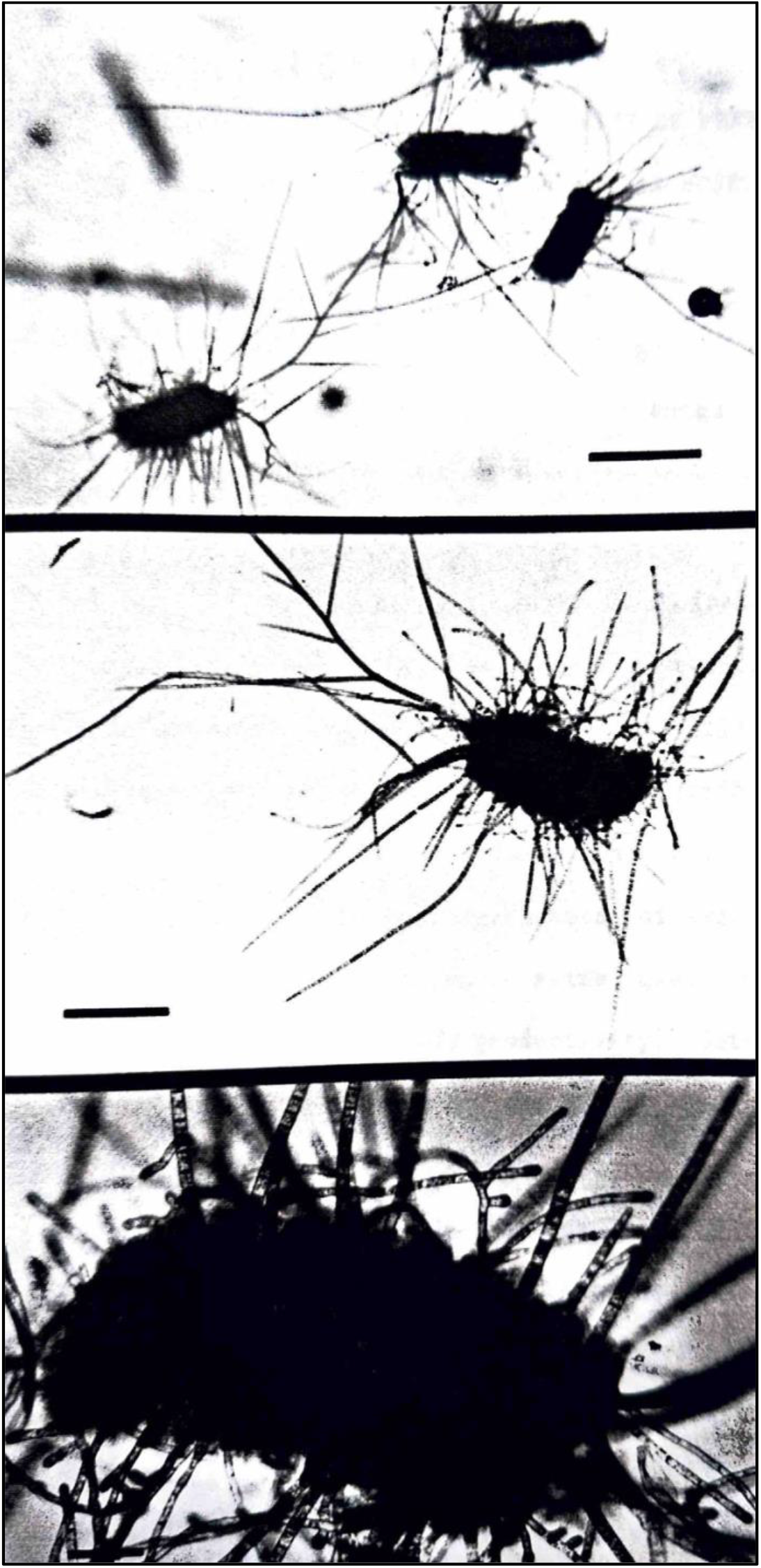
*Gammarus tigrinis* fecal pellets produced in culture with unialgal *Pilayella littoralis*. Pellets are all 25-28 days old. Top; scale bar = 1 mm. Middle; increased magnification and under slight pressure beneath coverslip to highlight algal filaments, scale bar = 0.5 mm. Bottom; high magnification showing filaments developing from a fecal pellet. Scale bar = 0.1 mm.

The δ^13^C values for free-living *Pilayella littoralis* and *Gammarus tigrinis* overlapped in their ranges and averaged -17.7 o/oo and -17.4 o/oo, respectively. The isotope value of material collected from the *P. littoralis* community, but not cleaned of associated animals and other algae, was -17.6 o/oo. These values are not significantly different (t = 1.53, df = 5).

## Discussion

*Pilayella littoralis* survives ingestion by the amphipod *Gammarus tigrinis*. New filaments grow from fecal pellets produced by amphipods collected in laboratory culture and from the free-living algal community in Nahant Bay. Before these observations, fragmentation due to physical disturbance from wave action or by filament breakage following infection by the holocarpic chytrid (a fungus) *Eurychasma dicksonii* (Wright) Magnus were the primary mechanisms proposed to increase population numbers (Wilce et al. 1982). Survivorship after ingestion adds another means of reproduction. However, the relative contribution of the different fragmentation modes to population increases or maintenance is unknown.

The condition of the material in the guts of *Gammarus tigrinis* varied from unidentifiable debris to intact pieces. Discoid plastids of *Pilayella littoralis* could sometimes be discerned, even from material collected in the lower third of the amphipod gut. The implications of resistance to digestion for *P. littoralis* are magnified if one considers that other animals (isopods, small decapod crustaceans, and fishes) could similarly increase the rate of vegetative reproduction of free-living *P. littoralis*. Although the production of algal filaments from fecal pellets collected from field amphipods was low (less than 4 percent), this does not diminish the potential significance of the phenomenon since the amphipod population produces large numbers of pellets. Unlike some plant seeds that appear pre-adapted to survive ingestion with impenetrable outer coverings, how *P. littoralis* survives ingestion by *G. tigrinis* is unknown. Rapid transit through the amphipod gut or inefficient digestion are likely explanations. The phenomenon may be widespread as *P. littoralis* also survived ingestion by two limpets (Santelices and Correa 1985) and a closely related algal genus, *Ectocarpus* sp., survived ingestion by seven herbivores (Santelices and Correa 1985, Santelices and Ugarte 1987).

Several additional observations are noteworthy. First, we did not observe fecal pellets in collections of free-living *Pilayella littoralis*, but the pellets are negatively buoyant and could easily fall to the bottom. Second, cultures of unialgal free-living *P. littoralis* with *Gammarus tigrinis* produced abundant algal filaments less than 0.5 mm long. Notably, the size and number of filaments were not seen in unialgal cultures without amphipods. Regeneration of entire plants from filaments of this size occurred frequently in these cultures. Therefore, filament fragmentation by amphipod grazing should be considered another means of vegetative reproduction. Third, amphipods in culture were observed to graze the outer surface of fecal pellets, though we could not be sure what the amphipods were grazing. Despite this behavior, we observed algal germination from fecal pellets. Finally, while we cannot be certain that we observed coprophagy in our cultures, the phenomenon is common among marine amphipods (Shillaker and Moore 1987). The extent to which coprophagy occurs and its effects on algal germination rates from pellets produced by *G. tigrinis* in the field is unknown but seems unlikely to be important.

At the onset of our study, whether *Gammarus tigrinis* ate free-living *Pilayella littoralis* or only used the community as habitat was unknown. Since the use of stable isotopes is most effective in systems where primary production is limited to one or two isotopically distinct carbon sources (Kitting et al. 1984, Simensted and Vissar 1985, Suchaneck et al. 1985, Dunton and Schell 1987), this study presented an ideal opportunity to determine if *P. littoralis* carbon is assimilated by invertebrate consumers. The isotopic data presented here show that the amphipod *G. tigrinis* derives most of its dietary carbon from *P. littoralis*. Although other algae likely possess an isotopic signature similar to *P. littoralis*, their biomass in the floating algal drifts was small, and they are not common in amphipod guts. Other algae in the drift community include epiphytes of *P. littoralis* that occur seasonally (Wilce et al. 1982), diatoms, the green alga *Entocladia wittrockii* (Wille) Kylin, and the brown alga *Hecatonema maculan*s (Coll.) Sauvag. Also, drift algae other than *P. littoralis* periodically comprise a portion of the free-living community but are always short-lived. Carbon isotope values of macroalgae are between 17–20 o/oo (Fry and Sherr 1985) or 14–29 o/oo (Stephenson et al. 1984). Based on the δ^13^C data, the amphipods appear to have acquired most of their carbon from *P. littoralis*. However, a seasonal study of amphipod guts and faunal associates must confirm whether amphipods eat *P. littoralis* year-round.

Considering the nuisance aspects of *Pilayella littoralis* in Nahant Bay (Wilce et al. 1982, Miller 1988), the survival of ingested material and fragments generated by grazing makes the algal-amphipod relationship a two-edged sword. Amphipods reduce biomass through feeding, but inefficient digestion of filaments and the production of fragments by grazing can increase numbers within the population.

## Acknowledgments

SLM dedicates this paper to RTW, who provided guidance, enthusiasm, and friendship during my doctoral research. Thanks to Kenneth Dunton for the δ^13^C analyses of algae and amphipods. I am grateful to Robert Vadas for his discussion and encouragement throughout my graduate career. Thanks to Charlie Yarish who shared culture techniques. Andrew Davis and Timothy Briggs helped with sample collections. RTW received financial support from the Metropolitan District Commission: Division of Parks, Engineering, and Construction. SLM received funding from the Woods Hole Trust Fund, University of Massachusetts.

## Notes

### Competing Interest Statement

The authors have declared no competing interest.

